# NovoTax: prokaryotic strain identification from mass spectrometry-based proteomics data

**DOI:** 10.64898/2026.04.02.715787

**Authors:** Dennis Svedberg, André Mateus

**Author notes:** Corresponding author: André Mateus.

## Abstract

**Summary:** Traditional mass spectrometry-based proteomics typically requires prior knowledge of sample composition to match spectra to peptides. Yet, novel de novo peptide sequencing approaches can provide peptide sequences to identify the organism. Here, we introduce an end-to-end pipeline (NovoTax) to identify the closest prokaryotic proteome directly from raw bottom-up proteomics data. The approach combines peptide sequencing tools with an optimized implementation of peptide searching through an extensive proteome database. On a benchmark dataset of species isolates, we identified the reported species and strain in the majority of the cases, and showed that in discordant cases NovoTax was likely correct. Interestingly, NovoTax was also able to identify contaminating species in samples. The algorithm also identified the most abundant organisms in bacterial communities. In summary, NovoTax provides strain level identification of microbial samples enabling the downstream use of traditional proteomics search engines for a deeper proteome analysis.

**Availability and implementation:** The open-source software is available on GitHub at https://github.com/mateuslab-prot/NovoTax

## 1. Introduction

The identification of prokaryotic species in biological samples is fundamental to microbiology, e.g., in clinical diagnostics or environmental monitoring. Traditionally, this is accomplished through 16S rRNA sequencing or whole-genome sequencing. However, when the goal is to measure protein abundance, this adds another experimental step.

Conventional proteomics workflows depend on a database with the protein sequences of the organisms present in the sample. Yet, determination of peptide sequence from mass spectra can be achieved through manual annotation, rule based heuristics (Taylor and Johnson, 2001), probabilistic scoring (Zhang, et al., 2012), graph based approaches (Gutenbrunner, et al., 2021), or deep learning methods (Jun, et al., 2025; Sanders, et al., 2025; Yilmaz, et al., 2024; Zhang, et al., 2025). These sequences can then be mapped back to proteomes to infer the taxa present in the sample (Holstein, et al., 2026; Vande Moortele, et al., 2025). For prokaryotic species, the Genome Taxonomy Database (GTDB) contains a large number of bacterial and archaeal genomes, and provides quality metrics and a consistent framework for taxonomy (Parks, et al., 2026).

Although tools exist for de novo sequencing, peptide-to-proteome mapping, and taxonomic inference, there is currently no solution that begins with raw mass spectrometry files and produces a sample-specific protein database ready for conventional search engines. Here, we introduce NovoTax, a modular pipeline that performs de novo sequencing of raw mass spectrometry data from both data-dependent and -independent acquisition schemes, and maps the resulting peptides iteratively to genomic databases using a taxonomy-informed search strategy that increases speed when querying the large GTDB. NovoTax identifies the strain from analyses of single-species, including revealing contaminants, and from microbial communities.

### 2. Implementation

NovoTax was built with modularity in mind, allowing users to either use the default workflow or provide their own intermediate results. It consists of three steps that are summarised below (**Figure 1A**).

**Figure 1.**
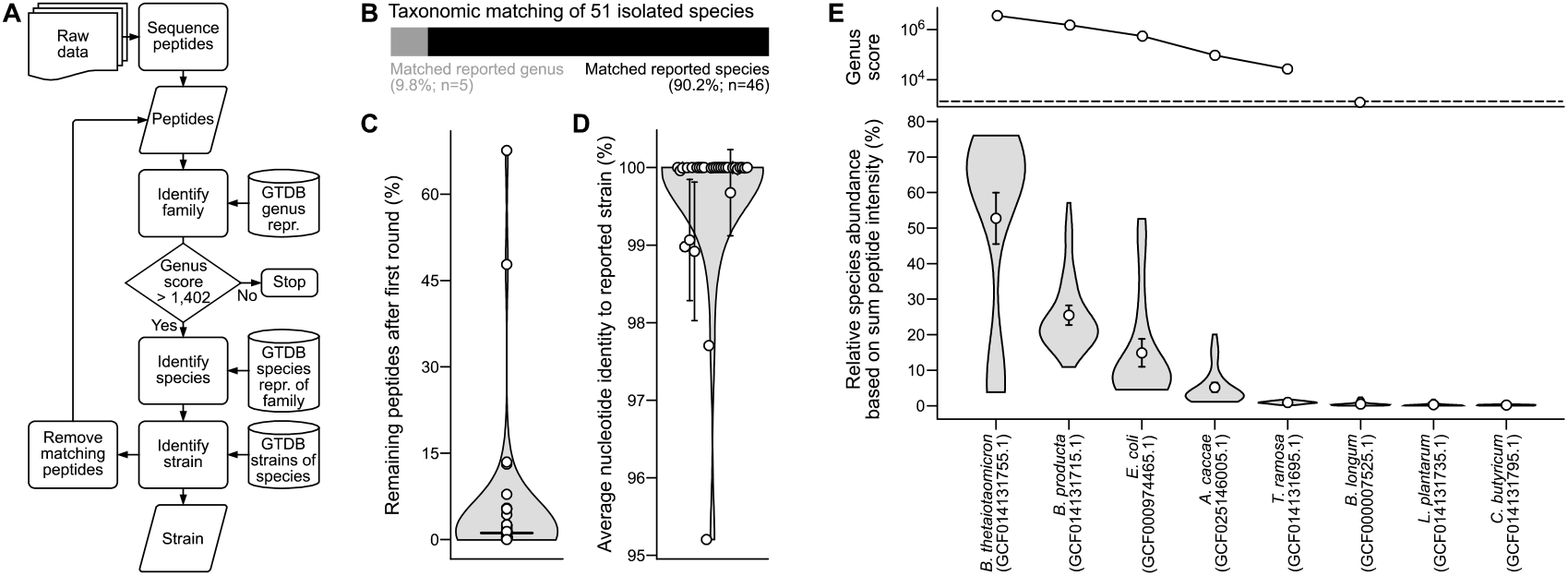
NovoTax identifies prokaryotic strains from raw mass spectrometry data. **(A)** NovoTax workflow that uses de novo predicted peptide sequences from raw mass spectrometry data to iteratively search GTDB. **(B)** Benchmarking of NovoTax using mass spectrometry files from single species proteomes. **(C)** Number of peptides that remain unassigned after matching the first strain for each raw file. **(D)** Average nucleotide identity between reported strain and the strain identified by NovoTax. **(E)** Species identified by NovoTax in a complex community and their relative abundance in the community based on the peptide identity.

### 2.1 De novo sequencing

NovoTax uses two default de novo sequencers, XuanjiNovo for data-dependent acquisition (DDA) and Cascadia for data-independent acquisition (DIA) data. The user points either to the files which they wish to sequence, or to a folder containing multiple files. The results are written to a peptide list including only peptides with a prediction confidence score above 0.8. If already available, the user can also provide a peptide table from another source, with optional scoring for further refinement.

### 2.2 Database peptide matching

Searches against GTDB (Parks, et al., 2026) are performed with MMseqs2 (Steinegger and Soding, 2017), by dividing them into three steps that minimize the size of the protein sequence database queried at any one time, speeding up queries, and lowering memory usage. The first search consists of a database including only genus representative species and, when no genus representative exists in GTDB for a family, a species with over 95% completeness and less than 1% contamination or the species with lowest contamination within the family. This reduces the database size from 732,475 proteomes (over 2.4 billion protein sequences) to 7,993 proteomes (about 24 million protein sequences). Once the genus is identified, a second search is performed against all species representatives for the family of that genus. In the third step, the peptides are searched against all strains for the identified species. In all steps, isoleucine is converted to leucine in both the list of queried peptides and in the database.

### 2.3 Taxonomy assignment

For each peptide, a score is calculated which is equal to the quality of the alignment (bitscore) divided by the number of proteomes matched, to penalize less informative peptides that match multiple proteomes. For each iterative step of the search, a sum of the matching peptide scores is calculated for each proteome (genus score), and the highest scoring proteome is selected. After the last search step, all peptides that are matched to the strain are removed and the non-matched peptides are searched again as described above. After the genus matching step, if the genus score is <1,402, the procedure is halted. This threshold was empirically chosen as described below.

## 3. Validation

To evaluate the performance of NovoTax, we used publicly available datasets (Lee, et al., 2022; Schape, et al., 2019; Wuyts, et al., 2023) that were retrieved from PRIDE (Perez-Riverol, et al., 2025).

### 3.1 Single species benchmark

Lee et al. (2022) collected a well curated dataset of 51 different bacterial species isolates consisting of 235 raw files (**Table S1A**). When running NovoTax on these files, we identified 46 species (90.2%) as the same as the reported species (**Figure 1B**). For the remaining 5 species, NovoTax identified members of the same genus (*Streptomyces griseoincarnatus, Lacticaseibacillus paracasei, Chryseobacterium rhizosphaerae, Agrobacterium tomkonis* and *Dorea formicigenerans*), but not the reported species (*S. griseorubens, L. casei, C. indologenes, A. tumefaciens* and *D. longicatena*). When performing a traditional proteomics search using MSFragger (Kong, et al., 2017) against a concatenated database of both the proteome identified by NovoTax and the proteome of the reported species, we observed that in all cases over 30% of peptides uniquely matched the proteome from NovoTax, while less than 2% matched the proteome of the reported species (**Figure S1; Table S1B**). This strongly suggested that the reported species was likely incorrectly unannotated.

In addition, for samples from two species (*Streptomyces venezuelae* and *Halomonas* sp. HL-93), we noticed that over 45% of the peptides from the de novo sequencer remained unassigned after the first iteration of the algorithm (**Figure 1C**). We suspected that these originated from another contaminant species in the same sample. Re-running the database matching step on the remaining peptides identified another species, and a traditional proteomics search against both proteomes identified over 39% of peptides from the potential contaminant in both cases (**Figure S2**). Based on this, we set an empirical threshold for NovoTax to continue running iteratively until the genus score reaches less than 1,402. This value is conservative and corresponds to the median genus score for the second round of the algorithm in these samples (**Figure S3**), i.e., we assumed that half the samples truly included only one species and remaining unassigned peptides correspond to poor predictions from the de novo sequencer.

For 32 species, Lee et al. (2022) reported a strain annotation that could be mapped back to GTDB. For 27 of them (84.4%), NovoTax identified a strain with over 99.5% average nucleotide identity (ANI) to the reported strain, i.e., the same or a very closely related strain (**Figure 1D**). For the strains that showed ANI <98%, performing a traditional proteomics search showed that the strain that NovoTax identified for *Cupriavidus necator* is likely a closer match to the analyzed strain, since a larger number of peptides uniquely matched that proteome (**Figure S4A**). While for *Delftia aciovorans* both the reported and NovoTax strains uniquely match a large fraction of peptides (>4%), which suggests that GTDB does not have a proteome for the exact strain that was analyzed (**Figure S4B**).

In summary, NovoTax can correctly identify the correct strain neighborhood for isolates of prokaryotic species provided that a proteome is available in GTDB. This includes identifying misannotated strains and even contaminating species.

### 3.2 Bacterial community benchmark

Next, we evaluated the performance of NovoTax on bacterial communities. We started with a simple community of eight species (Schape, et al., 2019). We collected all 90 raw mass spectrometry files and processed them together with NovoTax—we reasoned that this would allow capturing the largest number of species, since composition varied from sample to sample. While NovoTax identified only five of the eight species reported (**Table S1C**), these species represented on average over 99% of the peptides when performing a traditional proteomics search against a database with all the reported species (**Figure 1E**).

Finally, we tested the performance of NovoTax on a more complex community, by processing the raw files corresponding to the 96 h sampling time points from Wuyts et al. (2023), since the earlier samples were reported to be dominated by only two species. NovoTax identified 11 species in these samples, of which 9 were reported to be present among the most abundant species (**Figure S5**; **Table S1D**). Of note, these data were acquired in DIA mode, and thus we used Cascadia as a de novo sequencer (Sanders, et al., 2025). We noticed that a larger fraction of peptides could not be reliably assigned to any strain compared to DDA datasets, which could suggest that sequencing DIA raw files is less precise.

In conclusion, NovoTax is able to identify the most abundant members of a community.

## 4. Conclusions

NovoTax is an end-to-end pipeline to go from raw mass spectrometry-based proteomics data to taxonomic assignment of prokaryotic species. It enables retrieval of a protein fasta of the closest strain for downstream applications, which is important in prokaryotic species to correctly represent the gene content of the analyzed organism. The algorithm is fast enough to be run as a quality control for ordinary samples to ensure that the right strain is being analyzed and no contaminating organisms are present. It can also be used as a tool to discover microbial diversity in a community. It is packaged as modular Docker files and provides easy to interpret quality scores and taxonomic assignment, allowing non-experts to run it.

## Data availability

Benchmarking data was downloaded from the PRIDE repository (Perez-Riverol, et al., 2025) from datasets: PXD010000, PXD017035, PXD036445.

## Supplementary material

**Table S1**. List of processed raw files and results from NovoTax.

**Figure S1**. Proteomics search against concatenated databases of two proteomes: species reported to be present in the sample and species identified by NovoTax. Numbers refer to the number of unique and shared peptides.

**Figure S2**. Proteomics search against concatenated databases of two proteomes: species reported to be present in the sample and contaminating species identified by NovoTax.

**Figure S3**. Distribution of genus scores of remaining peptides after the first round.

**Figure S4**. Proteomics search against concatenated databases of two proteomes: strain reported to be present in the sample and strain identified by NovoTax. Numbers refer to the number of unique and shared peptides.

**Figure S5**. Species identified by NovoTax in a complex community and their relative intensity in the community based on 16S amplicon sequencing.

## Author contributions

D.S. and A.M. designed the study, developed and implemented the algorithm, and wrote the manuscript. A.M. acquired funding and supervised the study.

