## Supplementary figures and images for "NovoTax: prokaryotic strain identification from mass spectrometry-based proteomics data"

### Figure S1

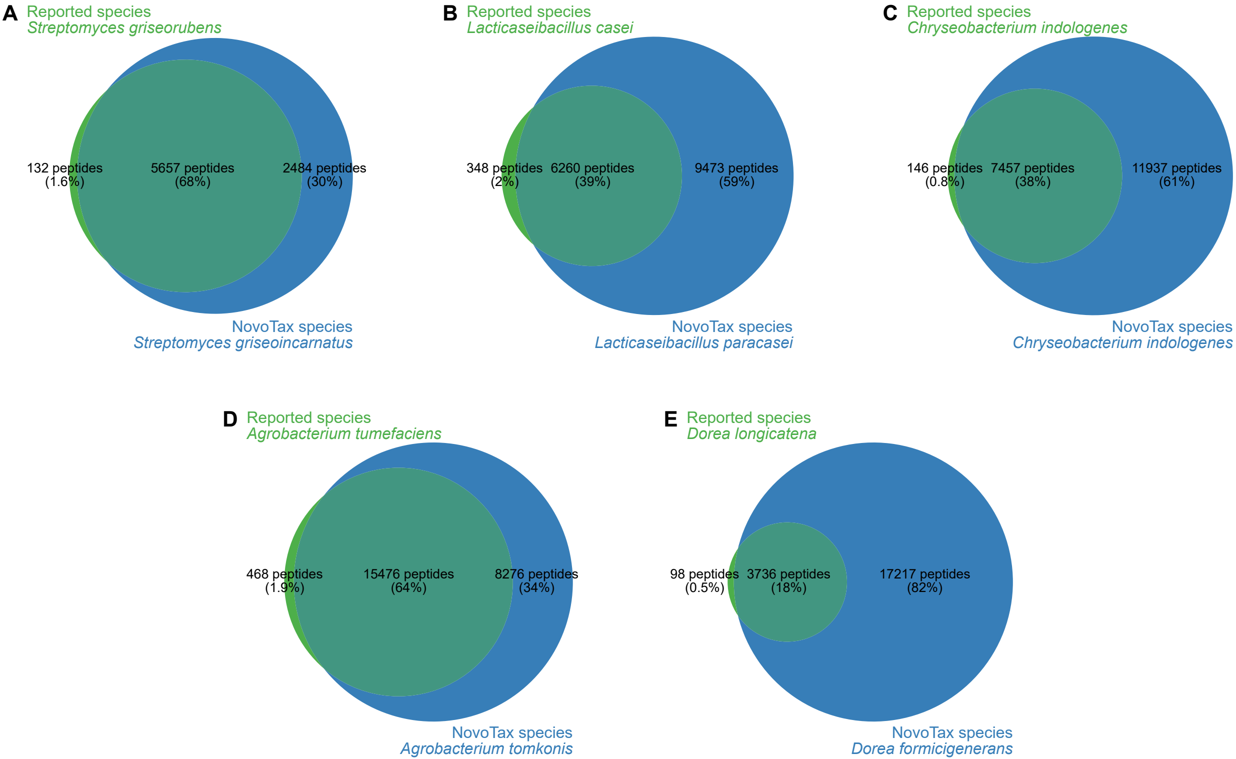

### Figure S2

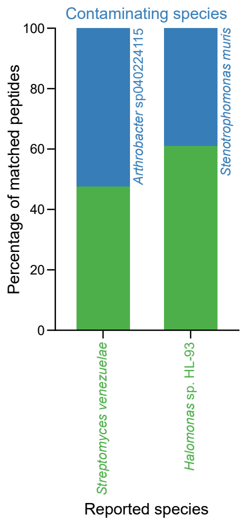

### Figure S3

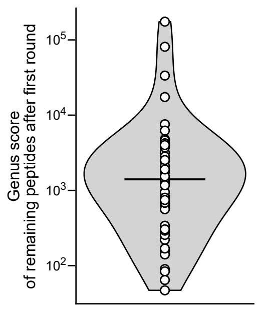

### Figure S4

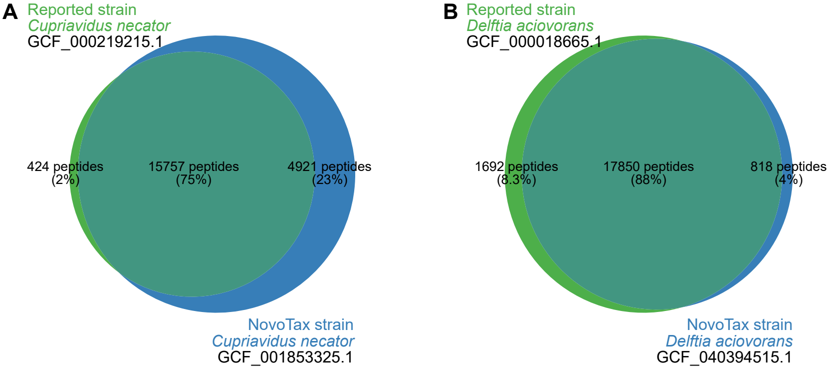

### Figure S5

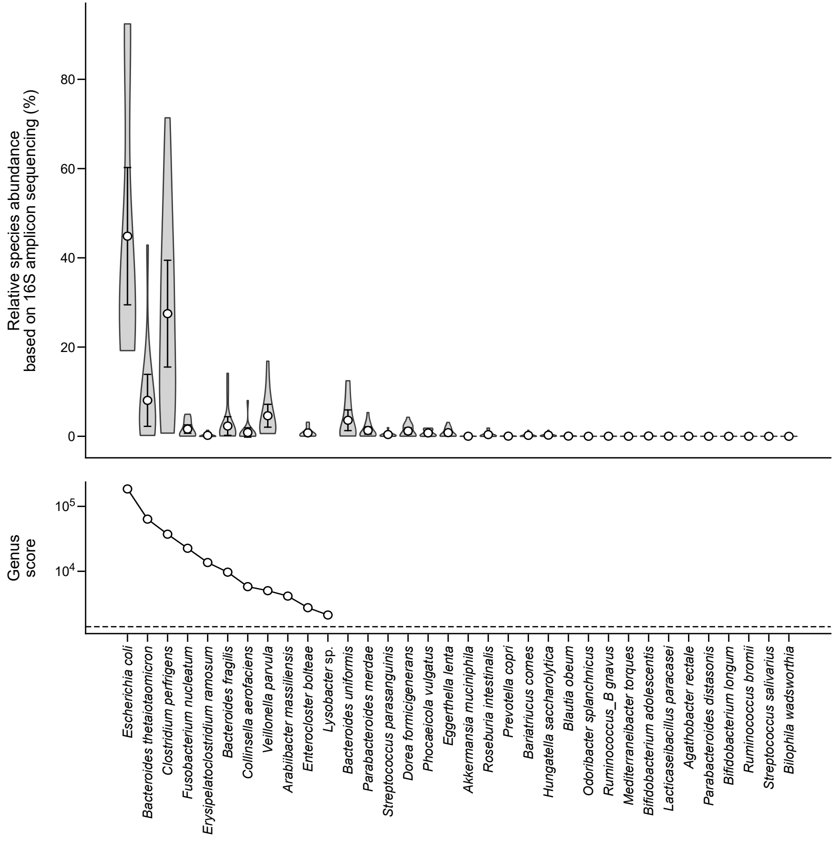
